# Probing Conformational Landscapes and Mechanisms of Allosteric Communication in the Functional States of the ABL Kinase Domain Using Multiscale Simulations and Network-Based Mutational Profiling of Allosteric Residue Potentials

**DOI:** 10.1101/2022.11.29.518410

**Authors:** Keerthi Krishnan, Hao Tian, Peng Tao, Gennady M. Verkhivker

**Affiliations:** Keck Center for Science and Engineering, Graduate Program in Computational and Data Sciences, Schmid College of Science and Technology, Chapman University, Orange, CA 92866, United States of America; Department of Biomedical and Pharmaceutical Sciences, Chapman University School of Pharmacy, Irvine, CA 92618, United States of America; Department of Chemistry, Center for Research Computing, Center for Drug Discovery, Design, and Delivery (CD4), Southern Methodist University, Dallas, TX 75205, United States of America

## Abstract

In the current study, multiscale simulation approaches and dynamic network methods are employed to examine the dynamic and energetic details of conformational landscapes and allosteric interactions in the ABL kinase domain that determine the kinase functions. Using a plethora of synergistic computational approaches, we elucidate how conformational transitions between the active and inactive ABL states can employ allosteric regulatory switches to modulate the intramolecular communication networks between the ATP site, the substrate binding region, and the allosteric binding pocket. A perturbation-based network approach that implements mutational profiling of allosteric residue propensities and communications in the ABL states is proposed. Consistent with the biophysical experiments, the results reveal functionally significant shifts of the allosteric interaction networks in which preferential communication paths between the ATP binding site and substrate regions in the active ABL state become suppressed in the closed inactive ABL form, which in turn features favorable allosteric couplings between the ATP site and the allosteric binding pocket. By integrating the results of atomistic simulations with dimensionality reduction methods and Markov state models we analyze the mechanistic role of the macrostates and characterize kinetic transitions between the ABL conformational states. Using network-based mutational scanning of allosteric residue propensities, this study provides a comprehensive computational analysis of the long-range communications in the ABL kinase domain and identifies conserved regulatory hotspots that modulate kinase activity and allosteric cross-talk between the allosteric pocket, ATP binding site and substrate binding regions.

## Introduction

Allosteric molecular events involve complex multiscale cascades of thermodynamic and dynamic changes that occur on different spatial and temporal levels.^1–7^ The human protein kinases that orchestrate functional processes in cellular networks are recognized as dynamic regulatory machines that exploit allosteric mechanisms for dynamic switching between the inactive and active kinase forms, thus enabling an efficient control of activity and adaptability during processes of signal transduction and catalysis.^8–13^ The wealth of structural knowledge about conformational states of the kinase catalytic domain, regulatory assemblies and ligand-kinase complexes is enormous and has dramatically advanced our understanding of the molecular determinants underlying kinase dynamics, function and binding.^11–13^ The dynamic equilibrium between the inactive and active kinase states can be affected and selectively modulated through activated mutations, by posttranslational modifications, and via ligand and protein binding events. This principle has been successfully exploited in discovery of small molecule kinase modulators.^14–,16^ Conformational transitions between kinase states are orchestrated by three conserved structural motifs in the catalytic domain: the αC-helix, the DFG-Asp motif (DFG-Asp in, active; DFG-Asp out, inactive), and the activation loop (A-loop open, active; A-loop closed, inactive). The conserved His-Arg-Asp (HRD) motif in the catalytic loop and the DFG motif are coupled with the αC-helix to form conserved intramolecular networks termed regulatory spine (R-spine) and catalytic spine (C-spine). A universal mechanism for dynamically driven allosteric conformational transformations and activation of kinases can be mediated by coordinated signal transmission through ordered hydrophobic architecture of the assembled spine networks.^17^ NMR studies have been instrumental in deciphering the allosteric energy landscapes of protein kinases, showing that the dynamic equilibria between the inactive and active states can be regulated through ligand binding to generate positive allosteric cooperativity and long-range communication networks between the distal kinase domain regions.^18, 19^ These studies suggested that structure and dynamics of the kinase catalytic core and residues surrounding the active site may have coevolved to optimize the intramolecular allosteric communication. Crystallographic and biochemical studies have provided a molecular framework for understanding mechanisms of ABL regulation by unveiling structural organization of the regulatory complexes.^20, 21^

Despite the established view of kinases as dynamic regulatory machines, the atomistic characterization of the intrinsic allosteric motions and functionally relevant transient states is often lacking due to large conformational transformations and short-lived kinase intermediates involved in the kinetics of the allosteric shifts. A series of pioneering NMR studies has provided a detailed atomistic picture of allosteric regulation in the ABL kinase by showing how interacting signaling modules cooperate with the kinase catalytic domain to form a multilayered regulatory mechanism that exploits various allosteric switches at different sites of the regulatory assemblies.^22^ A further considerable breakthrough in our understanding of the intrinsic conformational landscape of the ABL kinase domain has been recently achieved in the NMR experiments by Kalodimos and colleagues by revealing atomistic details of the hidden conformational states and determining the thermodynamically stable ground state of the ABL in the active state (Figure 1(a)) and two inactive conformations.^23^ By using state-of-the-art NMR techniques, this study captured the inactive short-lived ABL conformations I1 and I2 (Figure 1(b,c)) that occur only 5% of the time and are quite different from each other in the critical regions of the A-loop and the regulatory αC- helix motif. The unprecedented atomistic characterization of the conformational ABL ensemble has shown that these states are intrinsically present in the isolated ligand-free kinase domain, and allosteric transitions between these conformations can be modulated by substrate and ligand binding, allosteric interactions, and mutations.^23^

**Figure 1.**
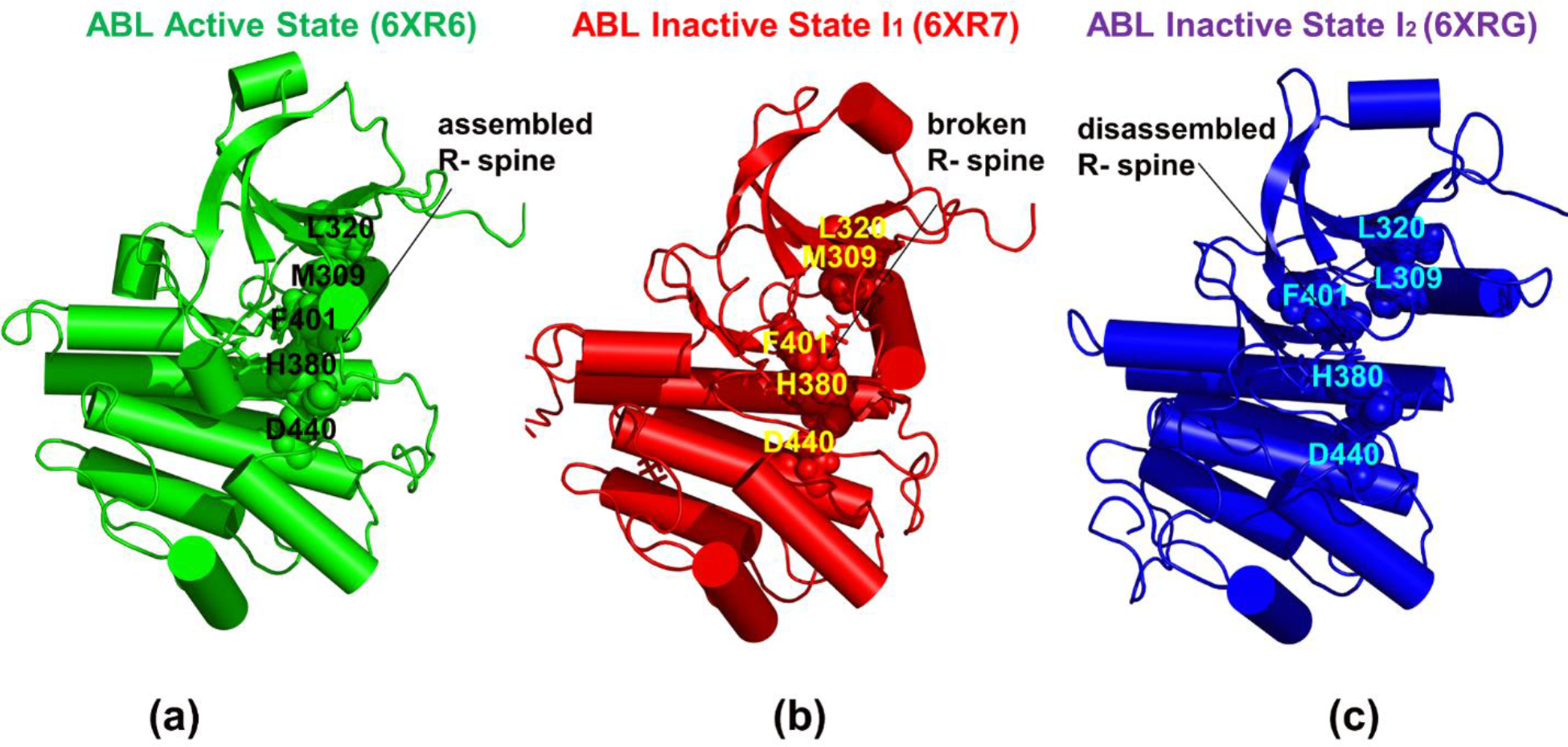
The NMR solution structure of the thermodynamically stable fully active ground state of the ABL kinase domain (pdb id 6XR6) is shown in green ribbons with the cylindrical helices (a). The NMR structure of ABL in the inactive state I1 (pdb id 6XR7) is in red ribbons (b) and the ABL structure in the closed inactive state I2 (pdb id 6XRG) is in blue ribbons (c). The R-spine residues M309, L320, H380, F401 and D440 in the active state and inactive state I1 are shown in green and blue spheres respectively. The R-spine positions in the inactive state I2 are L309, L320, H380, F401 and D440 and are shown in blue spheres. The structures point to similarities and differences in the key functional regions of the kinase domain exemplified by the αC helix, the A-loop, and the P-loop. In particular, A-loop in the inactive state I2 (c) adopts a completely different closed conformation. A fully assembled R-spine in the active ABL state (a) becomes partially broken in the inactive state I1 (b) and is fully disassembled in the closed inactive state I2 (c).

X-ray crystallography and hydrogen exchange mass spectrometry studies have characterized allosteric myristic pocket in the C-terminal lobe of the kinase domain showing that allosteric inhibitors GNF-2/GNF-5 and Asciminib can bind to a bent conformation of the αI helix that serves as a switch promoting the interactions with the SH2 domain in the regulatory complex and stabilization of the closed inhibited conformation.^24, 25^ At the same time, ligands that bind to the allosteric pocket but do not cause conformational change in the often αI helix have been found to often function as allosteric activators of kinase function.^26^ Remarkably, NMR spectroscopy studies have recently shown that Imatinib, that inhibits the catalytic site by inducing a specific inactive ABL conformation, can also act as an allosteric activator by binding to the myristic pocket and unexpectedly promoting kinase activity through binding competition between the ATP site an allosteric binding site, which becomes functionally relevant in the Imatinib-resistant ABL variants.^27^ Some of the latest NMR analyses suggested an alternative mechanism in which the disassembly of the inhibitory ABL-SH2-SH3 state and the opening of the regulatory core may be directly caused by Imatinib binding to the ATP site that allosterically enacts breakage of the inter-domain interfaces.^28–30^ These crystallographic and NMR studies have revealed functional role of allosteric interaction networks in the protein kinases, particularly highlighting the complexity of allosteric transformations and long-range communication networks between the ATP binding site, the substate binding region, and the allosteric site.

Computer simulations have explored the free energy landscape of ABL kinase suggesting that meta-stable ABL states are often structurally similar to known crystal structures of other kinases in complexes with a variety of inhibitors.^31–34^ Integration of molecular simulations and dynamic network modeling approaches described conserved allosteric pathways of the ABL and EGFR kinase regulation, providing a mechanistic model of allosteric communications between the ATP-binding and regulatory sites.^35–38^ Atomistic modeling of the ABL regulatory complexes bound to allosteric inhibitors and activators has shown that these small molecules can differentially modulate protein kinase activities by altering allosteric communications between the allosteric pocket and the binding regions.^39, 40^

A substantial challenge in investigating the allosteric mechanisms for large multi-domain kinase assemblies is the inherent difficulty of adapting experimental and computational methodologies to capture the intrinsic flexibility of these structures essential for functionality. Multidisciplinary structural biology studies that exploited synergies between NMR technologies, biophysical approaches and multi-scale computational methods have been particularly fruitful in uncovering the invisible dynamic aspects of diverse protein functions. In particular, this powerful toolkit of structure-centric techniques has been applied for mapping allosteric protein landscapes^41–44^, dissecting ligand-induced modulation of allosteric activation^45^, and characterization of allosteric communication networks that drive protein regulatory responses.^46–48^ A general approach of quantifying mutational effects for multiple molecular phenotypes using a specific implementation of multidimensional deep mutational scanning enabled a comprehensive characterization of allosteric mutations for many proteins.^44^ Using a combination of triple-resonance NMR and computational network analysis, the allosteric effects of specific kinase mutations and communication paths between regulatory elements and catalytic sites can be characterized.^47^ Using a combination of NMR spectroscopy, isothermal titration calorimetry (ITC), small-angle X-ray scattering (SAXS), and MD simulations, the intramolecular network of communications in PKA-C were elucidated, suggesting that the mutation-induced network changes may compromise highly cooperative regulation process leading to disease progression.^48^ Integration of NMR spectroscopy and surface plasmon resonance revealed dynamic communication networks of residues linking the ligand-binding site to the activation interface in the glucocorticoid receptor and identified a specific motif acting as a ligand- and coregulator-dependent switch for transcriptional activation.^49^ Solution NMR experiments and Gaussian-accelerated molecular dynamics (GaMD) simulations examined the structural and dynamic determinants of allosteric signaling within the CRISPR-Cas9 HNH nuclease, advancing our understanding of the allosteric pathway of activation.^50^ A further integration of NMR with multi-microsecond molecular dynamics (MD) simulations and graph-based network modeling probed the effects of mutations on the structure and allosteric communication within the CRISPR-Cas9 system, showing that mutations responsible for increasing the specificity of Cas9 alter the allosteric structure of the catalytic HNH domain.^51^ NMR chemical shift covariance (CHESCA) and projection (CHESPA) analyses^52–55^ can identify residue interaction networks that show correlated changes in chemical shifts due to allosteric perturbations caused by ligand binding or mutations designed to modulate allosteric conformational equilibria. Using statistical comparative analyses of the NMR chemical shift variations elicited by the selected perturbations, CHESCA approach characterizes perturbation-specific chemical shift patterns serving as signatures of allosteric mechanisms.

In the current study, we propose and integrate several computational approaches to examine principles of the ABL kinase domain allostery where the predicted allosteric signatures can be directly compared with structural and biophysical experiments. Despite unique insights provided by structural approaches in characterization of the conformational landscapes of the ABL kinase, the dynamic and energetic details underlying conformational rearrangements in the allosteric networks and role of allosteric regulatory switches within each of these states are not fully established. MD simulations are combined with the distance fluctuation coupling analysis of the ABL conformational ensembles to examine allosteric role of the key functional regions of the kinase domain. We introduce a perturbation-based network approach that implements a “deep” mutational profiling of allosteric residue propensities and interactions in the ABL states. In this method, using conformational ensembles of the ABL structures, we perform systematic modifications of protein residues and evaluate their effect on allosteric interactions and intramolecular long-range communications in the protein structure. By combining MD simulations with dynamic network modeling methods, we show how conformational changes between the active and inactive ABL states can affect allosteric couplings and modulate the intramolecular communication networks between the ATP site, the substrate binding region, and the allosteric myristic pocket. We determine that the preferential communication paths between the ATP binding site and substrate regions in the active state become partly suppressed in the closed inactive state I2 where in turn allosteric communications between the ATP site and the allosteric binding pocket become prevalent. By integrating the results of MD simulations with dimensionality reduction methods and Markov state models (MSM) we analyze the kinetic transitions between the ABL conformational states and show that the ABL kinase domain can transition following the path from the active state to the inactive state I1 and then to the inactive state I2, while direct transitions from active state to inactive state I2 are not favorable. Atomistic characterization of the dynamic allosteric changes and identification of the allosteric regulatory hotspots provide novel insights into role of allostery in diverse kinase functions, particularly in a mechanism underlying long-range control over communications between the ATP binding site, the substrate binding region, and the allosteric binding site. The results of this study suggest that probing the intramolecular communication networks in the ABL conformations through targeted modifications of vulnerable network links may be useful for engineering and modulating kinase activities.

## Materials and Methods

### MD Simulations

The crystal structures of the three ABL conformations in the active state, the inactive state I1, and inactive state I2) were taken from the Protein Data Bank (PDB), (PDB ID 6XR6, 6XR7, and 6XRG, respectively).^56^ The structures were further pre-processed through the Protein Preparation Wizard (Schrödinger, LLC, New York, NY) and included the check of bond order, assignment and adjustment of ionization states, formation of disulphide bonds, removal of crystallographic water molecules and co-factors, capping of the termini, assignment of partial charges, and addition of possible missing atoms and side chains. A box of TIP3P water molecules was used to simulate each system. Assuming normal charge states of ionizable groups corresponding to pH = 7, sodium (Na+) and chloride (Cl-) counter-ions were added to achieve charge neutrality and a salt concentration of 0.15 M NaCl was maintained. All Na^+^ and Cl^-^ ions were placed at least 8 Å away from any protein atoms and from each other. The topology and coordinate files for MD simulations were prepared using tleap^57^ and all-atom MD simulations were performed with AMBER ff14SB force field.^58^ Energy minimization after addition of solvent and ions was conducted using the steepest descent method for 100,000 steps in each ABL state. In the preparation stage, 100 ps of NVT ensemble Langevin MD simulations were conducted, followed by 200 ps of isothermal– isobaric ensemble (NPT) simulations at 1 atm and 300 K. Three independent 1µs NVT MD simulations were conducted for each of the ABL states. All MD simulations were conducted using the GPU version of the OpenMM software environment.^59^ Long-range electrostatic interactions were calculated using the particle mesh Ewald method^60^ with a real space cut-off of 1.0 nm. SHAKE method was used to constrain all bonds associated with hydrogen atoms. Simulations were run using a leap-frog integrator with a 2 fs integration time step.

### Distance Fluctuations Coupling Analysis of Allosteric Residue Propensities

The stability and allosteric propensities of protein residues were evaluated using distance fluctuation coupling analysis of the conformational ensembles. The computations and analysis are rooted in the protein mechanics approach^61, 62^ in which the fluctuations of the mean distance between a given residue and all other residues in the conformational ensemble are converted into distance fluctuations coupling indexes (DFCI) that measure the energy cost of the residue deformation during MD simulations. Our previous studies^39, 63^ showed that the DFCI parameter can correlate with allosteric residue propensities/potentials and the mean square fluctuations between a pair of residues could provide an accurate estimate of the signal commute time. The communication propensity of a pair of residues is inversely related to their commute time CT(i, j) expressed as a function of the variance of the inter-residue distance :

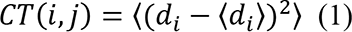

In our study, MD simulations of the ABL structures are analyzed by computing the fluctuations of the mean distance between each atom within a given residue and the atoms that belong to the remaining residues of the protein. The DFCI index can provide a measure of the allosteric potential for each residue and is calculated by averaging the distances between the residues over the simulation trajectory using the following expression:

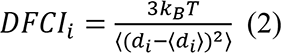

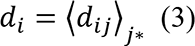

is the instantaneous distance between residue *i* and residue *j*, *k_B_* is the Boltzmann constant, *T* =300K. denotes an average taken over the MD simulation trajectory and *d_i_* =〈*di_j_*〉*_j_*_*_ is the average distance from residue *i* to all other atoms *j* in the protein (the sum over *j*_*_ implies the exclusion of the atoms that belong to the residue *i*). The reported DFCI index for each residue is computed as the average of the distance fluctuations for all its atoms *^i^ .* It should be noted that in addition we also computed mean fluctuations of a given residue by using *C_α_* atom positions as well as the reduced representations with a single pseudo-atom per residue^61, 62^ and a more refined model in which each amino acid is represented by one pseudo-atom located at the C*α* position, and either one or two pseudo-atoms representing the side chain.^64^ For clarity and consistency of the discussion, the reported DFCI indexes are based on full all-atom protein representation.

### Dimensionality Reduction Analysis of the Conformational Ensembles

MD equilibrium trajectories were analyzed using ivis dimensionality reduction method which is is a machine learning model that applies a Siamese neural network architecture composed of three identical neural networks.^65, 66^ Each network is composed of three dense layers with sizes of 500, 500, and 2000, followed by a final embedding layer of two neurons. The layers preceding the embedding layer use the SELU activation function:

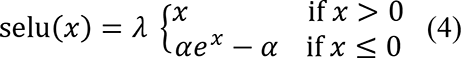

The values of *α* and *λ* are set as the default values of 1.6733 and 1.0507, respectively.^67^ The weights in these layers are initialized randomly using LeCun normal distribution. A linear activation is used in the final embedding layer initialized with Glorot’s uniform distribution.

During training, a triplet loss function is calculated as:

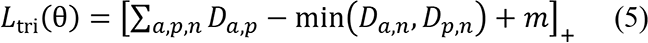

where *a, p*, and *n* are various kinds of points. *a*: point of interest, referred to as anchor point; *p*: positive points, selected through *k*-nearest neighbors (KNNs); *n*: negative points, randomly selected from the rest of the samples. A margin (*m*), the minimum distance gap, was set to the default value of 1. Euclidean distance (*D*) between data points is calculated to quantify the similarity:

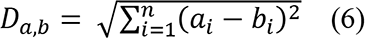

Adam optimizer with default learning rate was applied to train ivis model with GPU acceleration. ivis package (v 2.0.7) (https://pypi.org/project/ivis/) was used to implement the algorithm.

### Markov State Model

Stochastic Markov state models (MSMs)^68–70^ have become increasingly useful states-and-rates network models with the developed software infrastructure^71–74^ for describing the transitions between functional protein states and modeling of allosteric events. In MSM, protein dynamics is modeled as a kinetic process consisting of a series of Markovian transitions between different conformational states at discrete time intervals. A specific time interval, referred to as lag time, needs to be determined to construct transition matrix. MSM depends on the simulation distribution of the projected low dimensional space obtained by the ivis dimensionality reduction method. First, a MiniBatch *k*-means clustering method is conducted to assign each simulation frame to a microstate. Macrostates were clustered based on the Perron-cluster cluster analysis (PCCA)^75^ and are considered as kinetically separate equilibrium states. The transition matrix and transition probability were calculated to quantify the transition dynamics among macrostates.

The value of the lag time, as well as the number of macrostates, is selected based on the result of estimated relaxation timescale.^76^ The corresponding transition probability is calculated as:

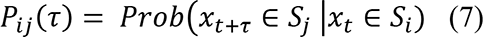

A proper lag time is required as the MSMs are required to be Markovian at the chosen lag time which can be assessed through the convergence of implied relaxation timescale. The implied timescale can be calculated using the eigenvalues (*λ*_*i*_) in the transition matrix as:

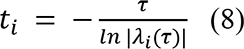

MSMs were constructed and the implied timescales were calculated using the PyEMMA package.^77^ Based on the transition matrix we obtain implied timescales for transitioning between various regions of phase space and use this information determines the number of metastable states. The number of metastable states also defines the resolution of the model by determining how large a barrier must be in order to divide phase space into multiple states.

### Dynamic Network Analysis

A graph-based representation of protein structures^78–80^ is used to represent residues as network nodes and the inter-residue edges to describe non-covalent residue interactions. We constructed the residue interaction networks using both dynamic correlations^81, 82^ and coevolutionary residue couplings.^83, 84^ To characterize allosteric couplings of the protein residues and account for a cumulative effect of dynamic and coevolutionary correlations, we employed the generalized correlation coefficient.^85^ The RING program was also employed for the initial generation of residue interaction networks.^86^ The ensemble of shortest paths is determined from matrix of communication distances by the Floyd-Warshall algorithm.^87^ Network graph calculations were performed using the python package NetworkX. The betweenness of residue *i* is defined as the sum of the fraction of shortest paths between all pairs of residues that pass through residue *i* :

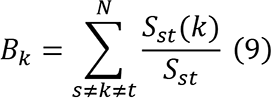

where *S*_*st*_ denotes the number of shortest geodesics paths connecting s and *t,* and *S*_*st*_(*k*) is the number of shortest paths between residues *s* and *t* passing through the node *k*.

The following Z-score is then calculated

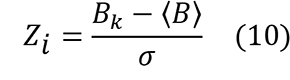

Through mutation-based perturbations of protein residues we compute changes in the average short path length (ASPL) averaged over all modifications in a given position. The change of ASPL upon mutational changes of each node is given as

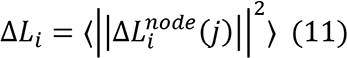

where *i* is a given site, *j* is a mutation and 〈 ⋯ 〉 denotes averaging over mutations. Δ*L*^*node*^_i_(*j*) describes the change of ASPL upon mutation *j* in a residue node *i*. Δ*L*_*i*_ is the average change of ASPL triggered by mutational profiling of this position.

Z-score is then calculated for each node as follows:

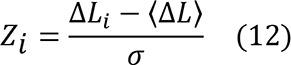

The ensemble-averaged Z –scores ASPL changes are computed from network analysis of the conformational ensembles using 1,000 snapshots of the simulation trajectory for the native protein system.

### eXtreme Gradient Boosting

eXtreme Gradient Boosting (XGBoost) is a powerful tree-based machine learning model^88^ with wide application in predicting protein structures and functions. During training, a new decision tree model is added iteratively. For a data *x*_*i*_, the final prediction is given by the sum of all decision tree models.

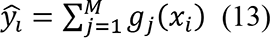

where *M* number of decision tr is the number of decision trees and *g*_*i*_ is a decision tree model.

The objective function in XGBoost is composed of a loss function *l* and a regularization term Ω to overcome overfitting:

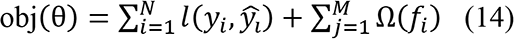

where 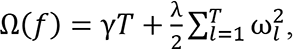, with *T* representing the number of leaves, and *γ, λ* being the regularization coefficients. The number and the maximum depth of tree models were fine-tuned and set as 300 and 4, respectively. The default values of other parameters were used. The XGBoost algorithm is implemented in xgboost package (version 1.6.2). (https://pypi.org/project/xgboost/)

## Results and Discussion

### MD Simulations of the ABL Kinase Allosteric States

We performed multiple independent all-atom MD simulations for the ABL kinase states using the structures of the isolated ABL kinase domain in its active conformational state (pdb id 6XR6) and two inactive conformational states I1 and I2 (pdb id 6XR7, 6XRG). All these states have been shown to be intrinsic to the unbound ABL kinase domain and allosteric structural transitions between these major conformational states could drive the kinase activities and response to the substrate and ligand binding (Figure 1).^23^ Structural analysis showed that the inactive ABL states (Figure 1 (b,c)) are very different, in which the regulatory DFG motif adopts distinct “out” conformations in the I1 and I2 states, while the A-loop and the αC helix undergo significant rearrangements. In particular, the αC helix moves from its active “αC-in” position to the intermediate position in the I1 state (Figure 1(b)) and to the “αC-out” inactive position in the I2 state (Figure 1(c)). In the active “αC-in” state a conserved αC helix residue E305 forms an ion pair with K290 in the β3 strand that coordinates the α and β phosphates of the ATP (Figure 1(a)). A substantial deviation from this arrangement is seen in the inactive states, which becomes particularly evident in the αC-out” I2 state where this hydrogen bond is largely broken (Figure 1(c)). The R-spine in ABL consists of M309 from the C-terminal end of the αC-helix, L320 from the β4-strand, F01 of the DFG motif in the beginning of the A-loop, H380 of the HRD motif in the catalytic loop, and D440 of the αF-helix (Figure 1(a,b)). The R-spine subnetwork is fully assembled in the active ABL kinase (Figure 1(a)) but becomes partially decoupled in the inactive state I1 (Figure 1(b)) and fully disassembled in the inactive state I2 (Figure 1(c)). The C-spine is comprised of hydrophobic residues (V275, A288, L342, C388, L389, V336, S457, and I451) connects the kinase lobes anchoring catalytically important sites to the C-terminus of the αF-helix.

MD simulations revealed important differences in the conformational dynamics of the ABL kinase core and long-range dynamic coupling between the binding sites that may be relevant for kinase inhibition and activation (Figure 2). By monitoring the root mean square fluctuations (RMSF) profile, we observed that the active ABL kinase state is more stable than the both inactive forms, displaying smaller thermal fluctuations in the intrinsically more flexible N-terminal lobe and A-loop (Figure 2 (a)). The dynamics of the inactive state I1 showed larger fluctuations but the overall shape of the RMSF profile remained mostly similar (Figure 2(a)). More significant dynamics differences were seen in the fully inactive state I2, with large thermal fluctuations observed in the middle of the A-loop that could obstruct the substrate Y412 position and inhibit substrate binding. Of particular importance are the dynamic behavior of the key regulatory elements of the catalytic kinase domain, particularly 400-DFG-402 motif and the R-spine residues.

**Figure 2.**
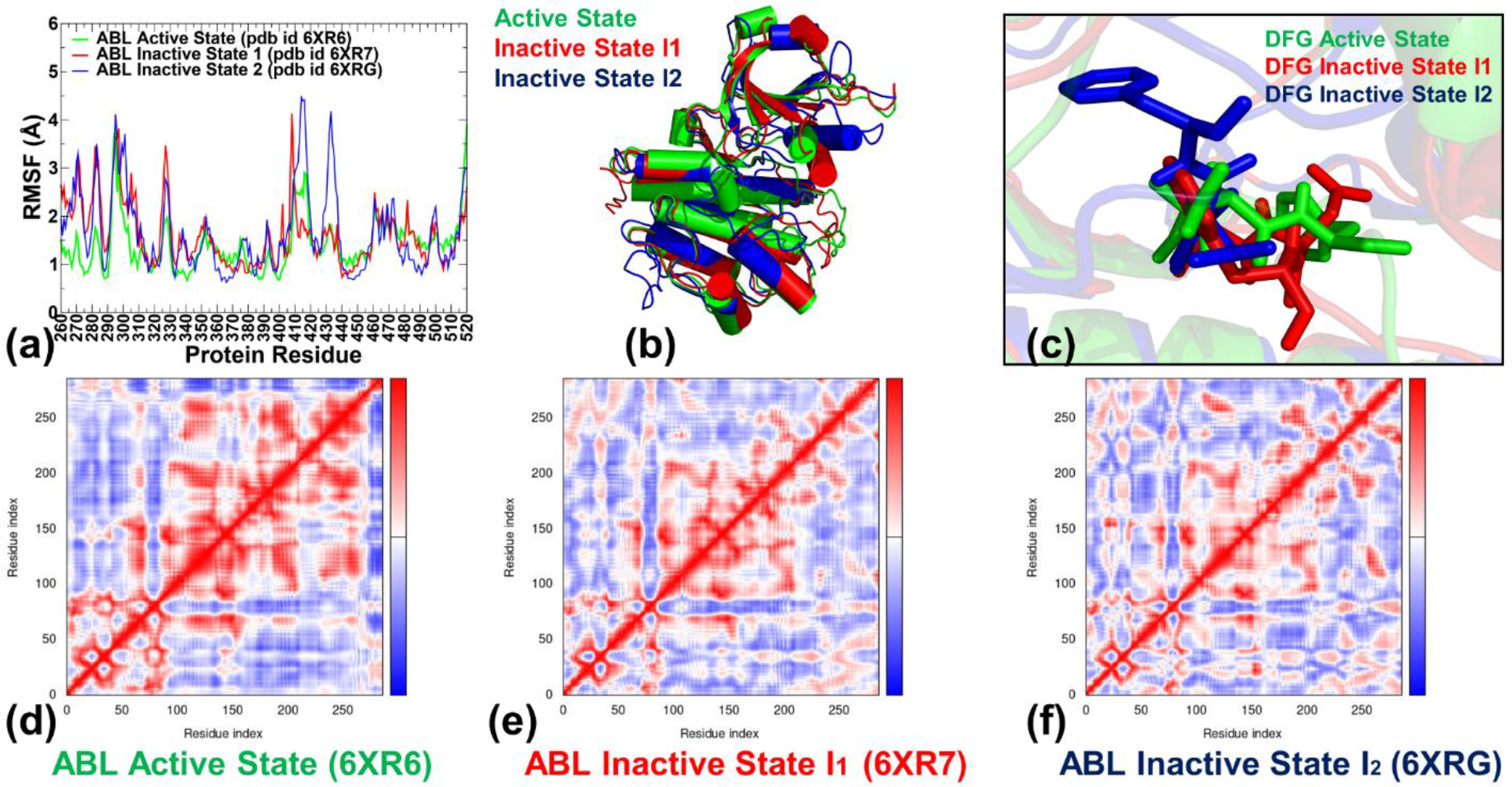
Conformational dynamics profiles of the ABL kinase domain states. (a) The root mean square fluctuations (RMSF) profiles are shown for the active ABL form in green lines, for the inactive I1 state in red lines, and for the inactive closed I2 state in blue lines. (b) Structural superposition of the ensemble-averaged conformations for the active state (in green ribbons), for the inactive I1 state (in red ribbons), and for the inactive I2 state (in blue ribbons). (c) Structural overlay of the regulatory 400-DFG-402 motif from the MD-averaged conformations of the active state (green sticks), the I1 state (red sticks) and the I2 state (blue sticks). A large movement of the F401 residue in the DFG-out conformation of the closed inactive I2 state can be seen. The covariance maps of dynamic cross-correlations between pairs of residues in the ABL active state (d), the inactive state I1 (e) and the inactive state I2 (f). Cross-correlations of residue-based fluctuations vary between +1 (correlated motion; fluctuation vectors in the same direction, colored in dark red) and -1 (anti-correlated motions; fluctuation vectors in the same direction, colored in dark blue). The values > 0.5 are colored in dark red and the lower bound in the color bar indicates the value of the most anti-correlated pairs.

The DFG-in motif in the active form flips by adopting the DFG-out conformation in both inactive states (Figure 2 (b, c)). This DFG movement is local in the I1 state but is much larger in the fully inactive state I2 where F401 translates by ∼10 Å to occupy a distinct hydrophobic pocket (Figure 2(c)). Notably, while experiencing pronounced structural rearrangements between the active and inactive ABL states, the DFG motif is stabilized in each of these ABL conformations and is anchored through a local interaction network formed with another indispensable stable element of the kinase core N387 residue. These residues displayed exceedingly small thermal fluctuations in both the active and inactive I2 state (Figure 2(a)) indicating that the interactions formed by the DFG motif are important for control of allosteric conformational changes in the ABL kinase. Indeed, the hydrogen bonding network between the N387 and D400 residues was experimentally shown to stabilize the DFG-in active conformation acting as a conformational switch for the DFG flips.^89^ The conformational fluctuations of the regulatory αC-helix (residues 299-311) were markedly smaller in the active ABL state where the stable “αC-in” conformation is coupled to the assembled R-Spine required for activation (Figures 2(a)). In the disassembled R-spine networks of the inactive states L320, H380, F401 and D440 residues experienced moderate fluctuations, while the larger displacements were seen for the αC-helix spine residue M309 (and L309 respectively in the I2) (Figure 2(a)). Interestingly, the thermal fluctuations of the regulatory αI-helix on the C-lobe (residues 504-517) are small in all three states, displaying even greater stability in the inactive ABL states. This helix is a key structural motif involved in formation of the allosteric binding pocket.^25–27^

The cross-correlation matrices of residue fluctuations along the low frequency modes highlighted broadly distributed and strong positive inter-residue couplings in the ABL active state (Figure 2(d)). In this form, long-range couplings were also observed between the kinase lobes, including the allosteric and ATP binding sites. Furthermore, the active ABL state featured strong positive couplings between the αF-helix anchoring the central core of the catalytic domain and the αD-helix and A-loop (Figure 2(d)). These correlations may reflect allosteric couplings along the C-spine and R-spine in the active state that link the ATP binding site and the substrate binding regions. These dynamic couplings are weakened in the inactive state I1 (Figure 2(e)) but the overall pattern of cross-correlations between different functional regions remained intact. A marked reduction of the positive long-range dynamic couplings was observed in the inactive state I2 (Figure 2(f)). In this state, a delocalization of coupled fluctuations in the catalytic core was observed, signaling not only a weakening of the long-range couplings but also a reorganization of the locally coupled structural elements in the kinase domain. In general, the cross-correlation maps of residue fluctuations in the catalytic domain were indicative of more cooperative allosteric interactions in the active kinase state as compared to looser and narrower patterns of correlated motions seen in the inactive states.

### Collective Dynamics Analysis of the ABL Conformational States Reveals Conserved Hinge Positions Controlling Modulation of Allosteric Changes

We characterized collective motions for the ABL states averaged over the 10 lowest frequency modes using principal component analysis (PCA) of the MD trajectories (Figure 3). The local minima along these profiles provide information about potential hinge regions that may control functional movements, while the maxima typically point to the protein regions undergoing global functional movements. It is assumed that functional movements along the pre-existing slow modes could drive allosteric transformations between the active and inactive states and provide insight into role of functional kinase residues in modulating population-shift allosteric changes between ABL states. The overall shape of the essential profiles and the key hinge centers were generally preserved across the three ABL states but showed considerable differences between globally moving regions in the inactive states (Figure 3). The predicted hinge positions 317- LVQLLGV-323 and 335-EFMTYGN-341 can mediate large-scale movements of the N-terminal lobe (Figure 3(a)). This finding is consistent with the experimental evidence^90^ suggesting that mutations in these regions can increase the hinge flexibility and modulate kinetics of allosteric changes between kinase states. According to the experimental studies, allosteric transitions related to the domain movement can be modulated by mutations into the hinge region to generate two mutants with increased flexibility.^90^

**Figure 3.**
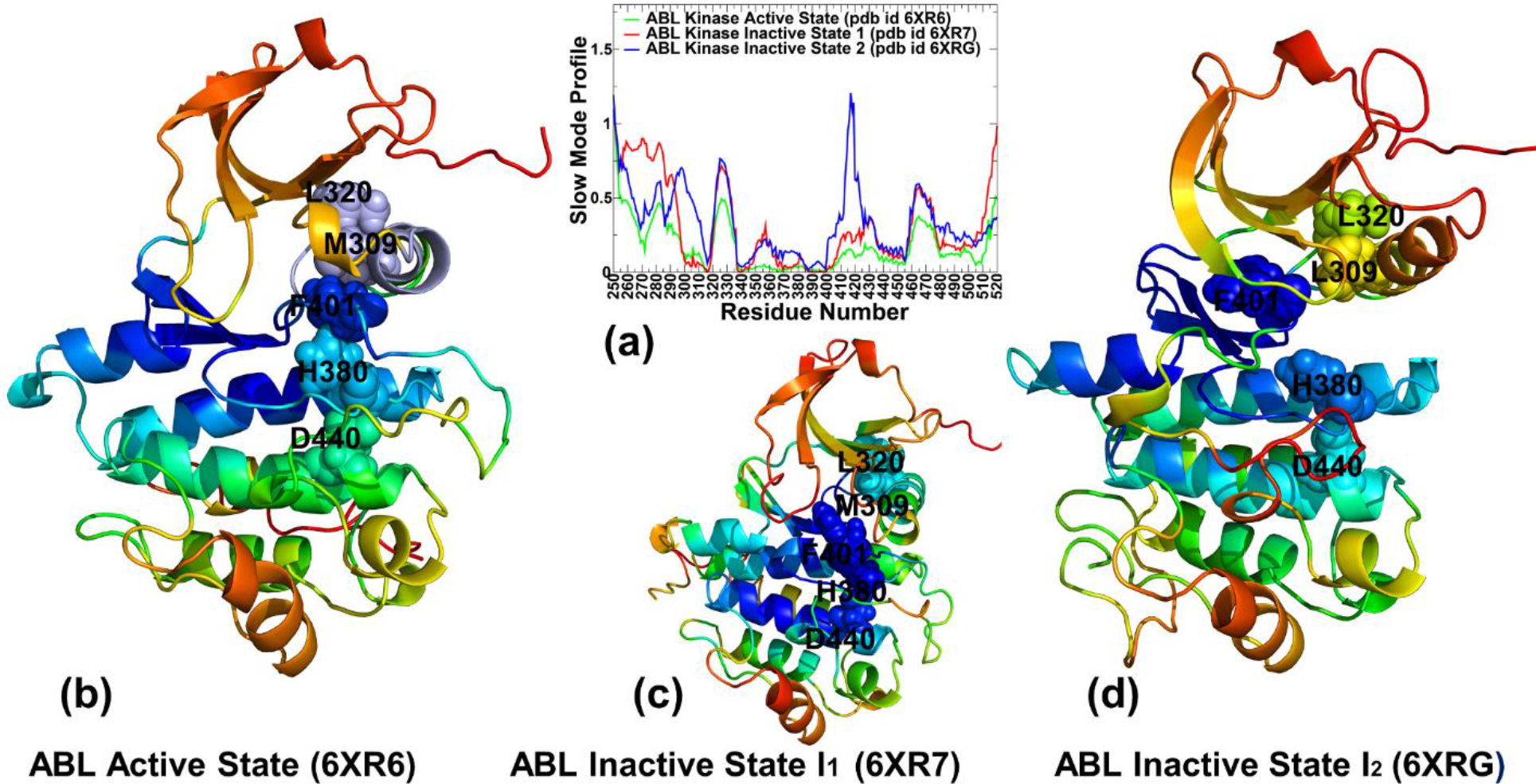
The collective dynamics profiles of the ABL conformational state. The essential mobility profiles are averaged over the first 10 major low frequency modes. (a) The slow mode profiles of the ABL kinase domain are shown for the active state (in green lines), the inactive I1 state (in red lines) and the inactive I2 state (in blue lines). Structural maps of the essential mobility profiles on PCA of the active state (b), the inactive I1 state (c), and the inactive I2 state (d). The mobility profiles are projected onto experimentally determined ABL structures shown ribbons and colored from blue to red according to the rigidity-to-flexibility scale determined by PCA. The R-spine residues are shown in spheres colored according to their level of rigidity/flexibility and annotated.

The importance of the hinge residues located between the N- and C-lobes of the kinase was also confirmed using NMR analysis which revealed that mutational disruption of the three molecular brake residues N549, E565, and K641 in FGFR2K (Q319, E335, K397 respectively in ABL) activate the enzyme.^91, 92^ These studies suggested that a functionally important allosteric pathway between the DFG-motif and the molecular brake residues is mediated by I547 (V317 in ABL) located away from the nucleotide and substrate binding pockets. Here, consistent with these studies, we found that dynamic changes in V317 are coupled with the F401 of the DFG motif (Figure 3). In addition, our analysis revealed another hinge cluster of locally interacting residues that is formed by residues N387, C388, A399 and the regulatory 400-DFG-402 motif (Figure 3(a)). According to our findings, the local interaction clusters of hinge residues anchored by V317 and N387 are dynamically coupled to the F401 of the DFG motif. Given the regulatory role of hinge centers in modulating allosteric transitions, we suggest that local structural perturbations of residues in these clusters may induce global rearrangements between the functional ABL states. The differences between the slow mode profiles of the ABL states are manifested in the distributions of the moving regions. The P-loop region (residues 267-275) in the N-terminal lobe displayed smaller functional displacements in the active state and the inactive I2 state, even though the P-loop adopts the kinked conformation in the active state and a more stretched conformation in the I2 state (Figure 3(a)). Notably, the shape of the slow mode profiles in this region featured a sharp local minimum (local hinge) that is precisely aligned with Y272 residue. Indeed, the packing interactions between Y272 and F302 and hydrogen bonding of Y472 with E305 are responsible for stabilizing P-loop and the αC helix-out conformation. At the same time, the P-loop in the inactive I1 state appears to exhibit large functional displacements in slow modes (Figure 3(a)). This may be indicative of the “transitional” nature of the inactive I1 state that acquires the increased mobility in the functional regions (including the P-loop) to facilitate adaptation of the fully inactive state. The slow mode profiles also pointed to similar and moderate displacements of the A-loop (residues 395-420) in the active state and inactive I1 states (Figure 3(a)) as the overall A-loop conformation is in an open conformation in both of these forms.

In contrast, significantly larger functional displacements can be afforded in the A-loop of the inactive state I2. We argue that functional movements of the P-loop and especially distorted A-loop in the inactive state I2 could provide driving force for allosteric changes and conformational adjustments of the ABL in the inactive state. Consistent with this assertion, Imatinib binding to the I2 state may induce additional structural changes in these regions required for optimizing the inhibitor-ABL interactions.^23^ Structural mapping of the slow mode profiles onto the ABL structures highlighted subtle but important variations in the distribution of the immobilized and globally flexible kinase segments in slow modes (Figure 3(b-d)). In the active state, the stable regions are aligned with the R-spine and C-spine regions linking the kinase lobes as well as the ATP active site and the substrate binding region (Figure 3(b)). The immobilized core becomes smaller in the state I1 (Figure 3(c)) where a fraction of the R-spine (H380, F401, and D440) is aligned with the hinge regions but the disassembled component in the αC-helix (M309, L320) appeared to be prone to larger displacements. A further readjustment of the moving regions could be seen in the state I2 (Figure 3(d)), in which only F401 residue of the R-spine remains rigid in the low frequency motions, while other components of the R-spine can undergo displacements thus compromising the integrity of the R-spine required for activating kinase function.

To summarize, the collective dynamics analysis revealed an important integrating role of the hinge cluster that includes locally interacting and strongly coupled stable residues N387, C388, A399, D400 and F401 that collectively may function as a global coordinator of the kinase functional movements. The dynamic couplings between this rigid hinge cluster and flexible functional regions of the αC-helix, P-loop and A-loop capable of undergoing coordinated movements can serve a potential mechanism for modulating allosteric transformations between the active and inactive ABL states.

### Distance Fluctuation Coupling Analysis of Stability and Allosteric Residue Preferences in the ABL Functional States

Using the results of MD simulations, we conducted the distance fluctuation coupling analysis and examined how these distributions and positions of the local peaks change in different ABL states (Figure 4). In this model, dynamically correlated residues whose effective distances fluctuate with low or moderate intensity are expected to communicate with higher efficiency than the residues that experience large fluctuations. By using the DFCI profile of the active state as the reference, we evaluated how dynamic changes in the inactive states can modulate the allocation of stable residues with high allosteric preferences (or allosteric potential) (Figure 4). A similar shape of the DFCI profile was observed for all ABL states, revealing that specific regions of the catalytic core could form hubs of dynamic couplings. To examine the role of these regions, we first analyzed the distribution obtained for the ABL active state. The multiple peaks distributed in the kinase domain reflected the presence of many spatially distributed allosteric centers mediating a broad network that links the kinase lobes and the binding sites (Figure 4(a)). Strikingly, the characteristic peaks of the distribution corresponded to the N387, C388 (C-spine), L389 (C-spine), A399, 400-DFG-402 residues, and W424 from the substrate binding region 24-WTAPE-428 of the C-lobe (Figure 4(a)). All these positions have been experimentally recognized as functionally important for kinase function, with the DFG motif being critical for differences between the inactive and active states, the C-spine residues responsible for catalysis, and the substrate binding residues. W424 is the experimentally validated allosteric node in which mutations (Y404A in protein kinase A) can decouple dynamic couplings and shift the dynamic equilibrium.^93^

**Figure 4.**
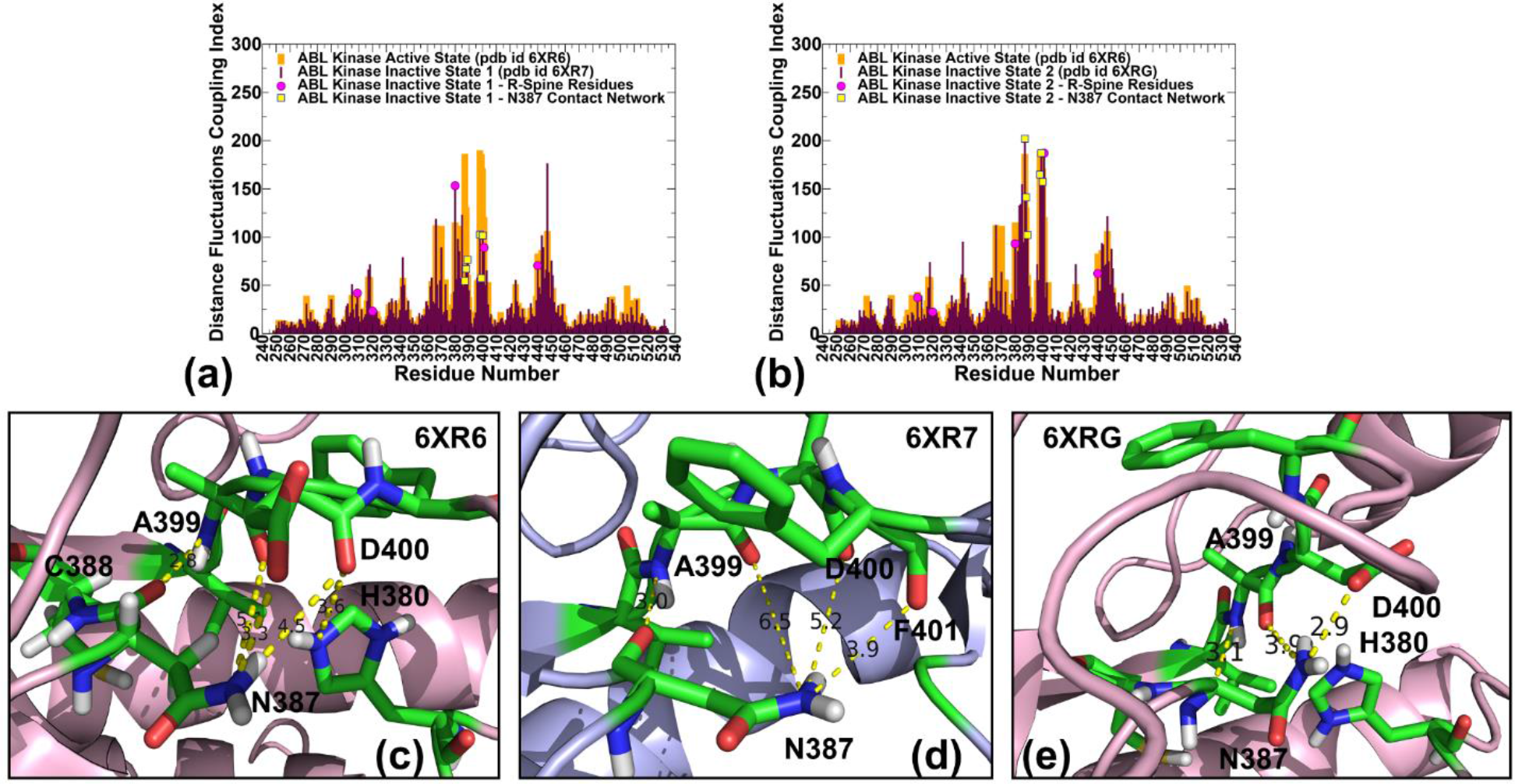
The residue-based DFCI profiles for the ABL conformational states. (a) The DFCI distributions for the active state (in orange bars) and the inactive state I1 (in maroon-colored bars). (b) The DFCI distributions for the active state (in orange bars) and the inactive state I2 (in maroon-colored bars). The positions of the R-spine residues (M309/L309, L320, H380, F401, D440) are shown in magenta-colored filled circles. The residues involved in the N387-mediated local contact network (N387, C388, A389, D400, F401) are shown in yellow-colored filled squares. A close-up of the local interaction cluster mediated by N387 in the ensemble-averaged conformation of the active form (c), the inactive I1 state (d), and the inactive I2 state (e). The interacting residues are shown in atom-colored sticks and specific interaction contacts are annotated.

In the inactive state I1, the DFCI distribution revealed an important change as the indexes for N387, A399, D400 and F401 were reduced. According to our model, this may indicate the increased flexibility and partial weakening of the interactions in this key hotspot cluster (Figure 4(b)). In the ABL inactive state I2 we observed a partial reorganization of the DFCI distribution peaks in which the depletion of some peaks may narrow the spectrum of mediating centers in the allosteric network, whereas the role of locally coupled N387, A399, D400, F401 and L403 residue is amplified as this cluster emerged as a single dominant hotspot of dynamic couplings in the kinase domain (Figure 4(b)).

The role of the predicted functional residues in mediating the dynamic inter-residue couplings in the ABL catalytic domain can be better understood by examining the local interaction clusters formed by these key sites in different ABL states. In the ABL active state, the carbonyl oxygen of N387 is hydrogen bound to NH of A399, and amide group of N387 makes several hydrogen bonds with the side chain and carbonyl oxygen of D400 (Figure 4(c)). A strong interaction cluster formed by N387 with A399 and D400 in the active state provides an additional support to the fully assembled R-spine and contributes to stabilization of the DFG-in active conformation. This is consistent with a critical role of N387 residue acting as a conformational switch of the ABL dynamics found in the functional studies showing that mutations of N387 that weaken the local interactions in this cluster can allosterically promote the increased global flexibility of ABL and facilitate faster Imatinib dissociation.^89^ Interestingly, a recent computational study showed that Imatinib can dissociate from the wild-type ABL and ABL-N387S mutant through two distinct pathways where the predominant Imatinib unbinding pathway through the kinase hinge region in the wild type ABL is altered in the ABL-N387 mutant and occurs via the *α*C-helix region.^94^ In the inactive I1 state the interactions mediated by N387 residue are weakened due to the loss of hydrogen bonding seen in the active state that could not be compensated by suboptimal N387 side chain contacts with the carbonyl oxygen of F401 (Figure 4(d)). On the other hand, in the inactive I2 state the hydrogen bonding mediated by the side-chain of N387 with the backbone of A399 and side-chain of D400 can be partly restored (Figure 4(e)). In addition, F401 of the DFG-out motif is rigidified in its inactive position through favorable contacts with F336 and L403 residues. Our results suggest that the structural changes in the local interactions mediated by N387 can be coupled with the potential allosteric role of these residues. This may partly explain why F401V mutation that enhances stability of the local interactions in this region could preferentially stabilize the inactive I2 state but not the I1 inactive form.^23^ In addition, the N387- mediated contact network in the I2 state can rigidify the H380-F401-D440 fragment of the distorted R-spine thereby disfavoring movements required to restore connections with M309/L320 N-terminal component of the R-spine (Figure 4 (e)). The observed reorganization of the intramolecular contacts between key stabilization centers could therefore promote a complete disintegration of the R-spine and alter the intramolecular allosteric network. Structural mapping of the major distribution peaks associated with the N387 interaction network that includes A399, D400 and F401 residues illustrated the coupling of this local cluster with the R-spine residues (Figure 5). In the active ABL state, the N387 cluster is tightly coupled to the fully assembled R-spine and together with Y412, R405 and R381 of the 380-HRD-382 motif form an interconnected stabilizing network connecting the kinase lobes and linking the binding sites (Figure 5(a)). The “grip” of the N387 mediating interactions on the partially disassembled R-spine in the inactive state I1 is reduced (Figure 5(b)). On the other hand, the N387 cluster could strengthen the disassembled R-spine in the inactive state I2 (Figure 5(c)).

**Figure 5.**
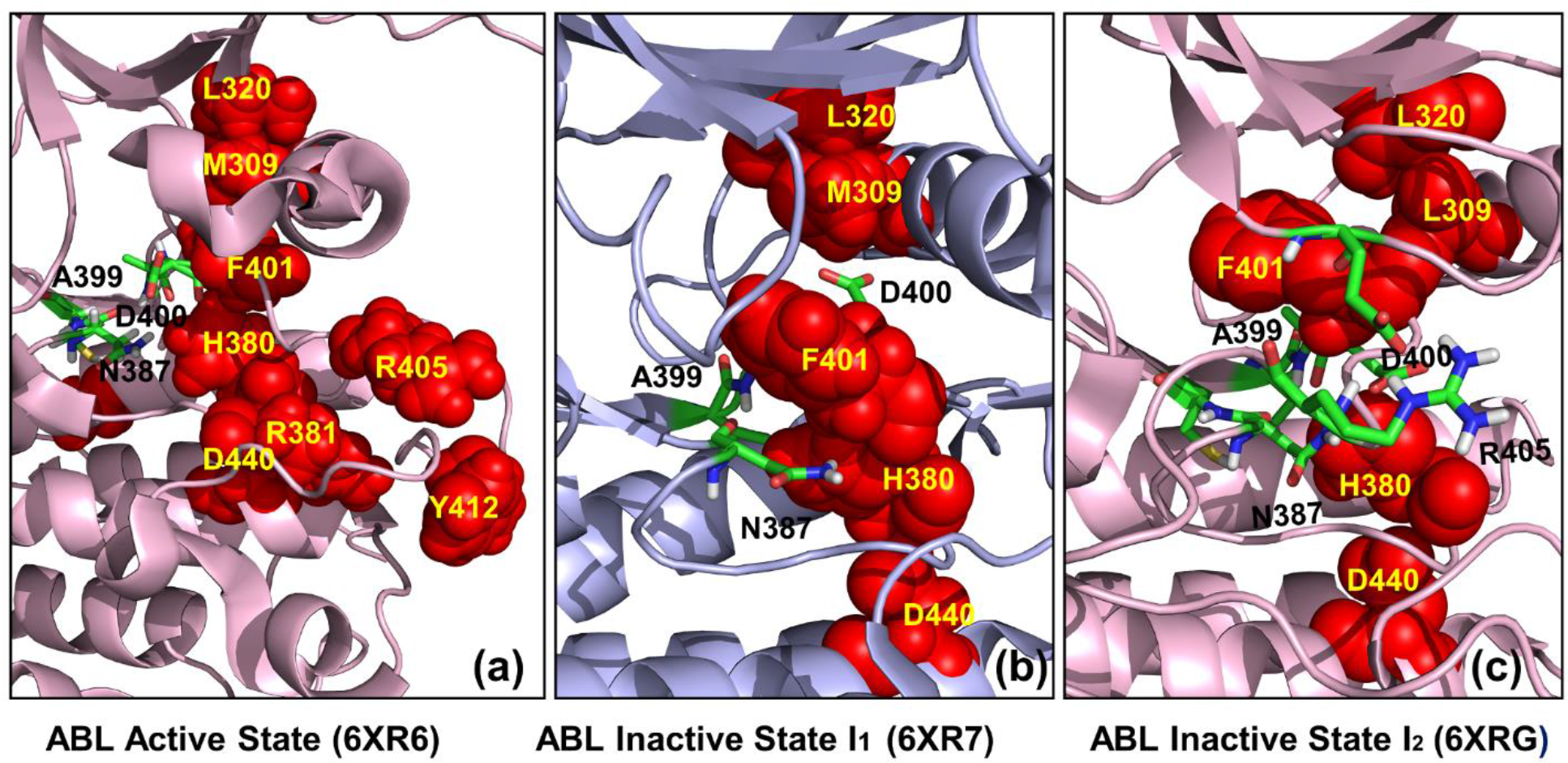
Structural mapping of the N387-medited local interaction cluster and the R-spine residues in the active state (a), the inactive I1 form (b), and the inactive I2 conformation (c). The residues involved in the N387-mediated local contact network (N387, A389, D400, F401) are shown in atom-colored sticks. The R-spine residues are shown in red spheres and annotated. The ABL structures are shown in light-pink colored ribbons.

To summarize, the DFCI analysis revealed a key role of locally interacting functional residues N387, A399, D400, and F401 that feature a high allosteric potential. Our results indicated that these allosteric centers together with the R-spine residues could modulate dynamic couplings and the intramolecular communication networks in the ABL states.

### Network-Based Mutational Profiling of Allostery : Conformation-Specific Reorganization of Allosteric Interaction Networks Modulates the Intramolecular Communications

The residue interaction networks in the ABL conformational states were built using a graph-based representation of protein structures^78–80^ in which residue nodes are interconnected through dynamic^81, 82^ and coevolutionary correlations.^83, 84^ In this method, we perform systematic modifications of protein residues and evaluate their effect on the dynamic inter-residue couplings and the residue interaction network organization as was initially proposed in our previous studies.^95^ By computing mutation-induced changes in the topological network parameters such as ASPL that characterize the efficiency and robustness of allosteric communications in the ABL states, we identify the allosteric regulatory hotspots as positions in which the average effect of mutations can cause a significant ASPL changes, leading to the network alterations that may compromise the efficiency of allosteric communications in the ABL states. The mutational profiling distribution in the active state featured a considerable number of peaks that are broadly distributed in the kinase domain (Figure 6(a,b)). The key sites that affect allosteric communication in the active ABL are precisely aligned with the R-spine residues L320, H380, F401 and D440 (Figure 6(a)). In addition, the local interaction cluster anchored by N387 is also involved in mediating the allosteric network in the active form. The distribution highlighted a noticeable peak in the substrate binding motif 424-WTAPE-428 of the C-lobe, which is consistent with the functional studies showing that W424 can act a conserved allosteric node linked to the R-spine residues.^93^ Structural mapping of the 20 residues featuring the highest allosteric potentials showed a broad and dense interaction network connecting the substrate site with the ATP site (Figure 6(b)). This network is determined by connectivity afforded through dynamic couplings between the αC- helix-in, the fully assembled R-spine and the substrate binding site in the active state. The predicted topological map of allosteric communications in the active ABL is consistent with the experimentally observed patterns of the intramolecular cooperativity in other kinases. For example, biochemical experiments identified a similar dynamically coupled allosteric network between the ATP- and substrate-binding sites in Src kinase, showing that the robust intramolecular communications and long-range cooperativity in the active state underlies the enzymatic kinase function.^96^ Our findings are also consistent with the NMR-based CHESCA analysis of correlation maps between functional network communities in the active state of PKA enzyme showing strong allosteric communications between the ATP-binding site, the αC-helix and the A-loop.^48^

**Figure 6.**
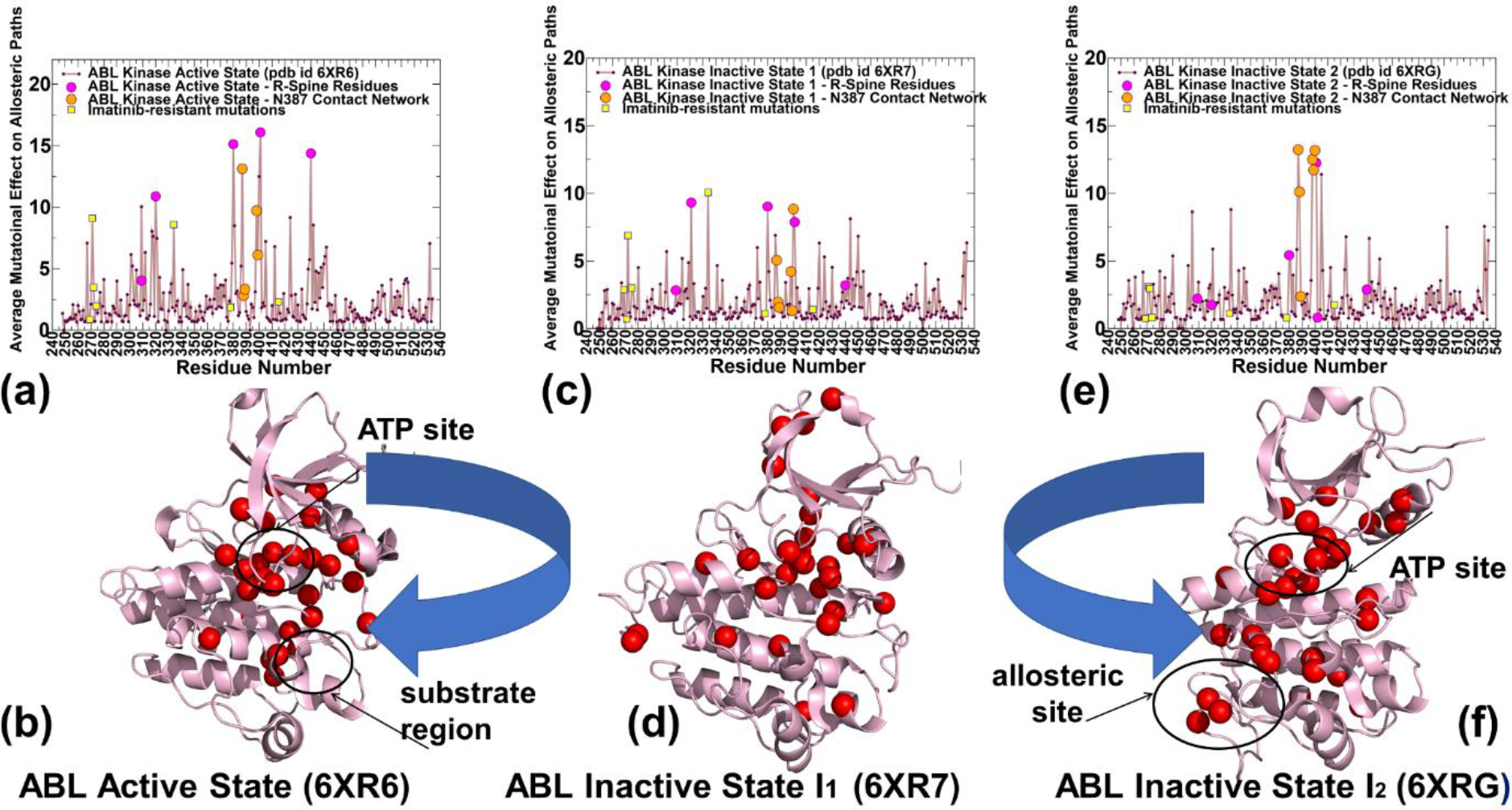
Mutational profiling of allosteric residue propensities in the ABL states. The residue-based Z-score profile estimates the average mutation-induced changes in the ASPL network parameter for the active state (a), the inactive I1 state (c), and the inactive I2 state (e). The profiles are shown in brown-colored lines. The positions of the R-spine residues on the distribution are highlighted in magenta-filled circles. The residues in the N387-anchored mediating local cluster (N387, C388, A399, D400, F401) are shown in orange-colored circles. The sites targeted by Imatinib-resistant mutations G269E, Q271H, Y272H, E274V, T334I, F378V and H415V are indicated by yellow-colored filles squares. Structural mapping of the top 20 kinase residues with the highest allosteric mediating potential in the active state (b), the inactive I1 state (d), and the inactive I2 state (f). The highlighted centers (shown in red spheres) are determined by the Z-score evaluation of the mutational effects on the intramolecular communication paths. The mapped allosteric mediating sites form the intramolecular communication routes connecting the ATP site, the substrate binding region and the allosteric pocket. The observed shifts in preferential communication paths between the active site and the inactive state I2 are highlighted.

We also found that modifications of residues from the allosteric binding pocket (R351, A356, L359, L360, A363, L448, L451, T453, Y454, M456 G482, C483, V487, F512, V525, L529) could induce only minor changes in the allosteric communications of the active form have a minor effect on the efficiency of the allosteric networks (Figure 6(a,b)). It appeared from this analysis that the long-range couplings between the allosteric site and the ATP site may be partially depleted in the active kinase form. These observations are consistent with recent biochemical and structural studies showing the lack of cooperation between ATP-competitive inhibitors that stabilize the αC- helix-in kinase conformation and allosteric inhibitors of ABL.^97–100^ At the same time, these studies suggested that conformation-selective ATP-site inhibitors that favor the inactive ABL conformations can afford a stronger binding cooperation of the allosteric site with the ATP site. An alternative view based on the NMR and isothermal titration calorimetry studies suggested that Asciminib can bind ABL concomitantly with both type-1 and type-2 ATP-competitive inhibitors to form ternary complexes.^98^

Mutational scanning of allosteric propensities in the inactive state I1 featured a similar shape of the distribution but revealed the reduced peaks associated with the R-spine positions M320, H380 and F401, while the peak corresponding to D440 completely vanished (Figure 6(c,d)). In the I1 conformation the αC helix remains in the “in” conformation with the catalytic bride K290-E305 intact as in the active state, but the DFG motifs swings into inactive “out” position rendering this state to be catalytically inactive.^23^ From structural mapping of the top allosterically-sensitive positions, it is evident that the allosteric network becomes weaker and involves only a portion of the R-spine. Nonetheless, our results suggested that allosteric communications between the ATP site and the substrate binding site in the inactive I1 state can be still functional but less robust than in the active form (Figure 6(d)). Interestingly, despite a partially disconnected R-spine, allosteric communication flow between the N-lobe and C-lobe of ABL can still be preserved owing to a large number of mediating centers in the functional regions. These observations are consistent with the NMR and network-based studies of allostery in p38γ kinases^47^, showing that a physically connected R-spine in the inactive state may be “dynamically” sufficient to enable allosteric signal propagation between the binding sites along the conserved architecture, while a completely assembled R-spine is a necessary prerequisite for phospho-transferase activity. Consistent with this study, we similarly found that stronger dynamic residue correlations and a broader interaction network in the active ABL form ensures efficient allosteric communication mediated by the fully assembled R-spine.

The structural and dynamic changes in the inactive I2 state appeared to incur more significant rearrangements in the interaction network in which contributions of the R-spine residues with the exception F401 are reduced (Figure 6(e)). The invariant cluster of allosteric hotspots includes N387, C388, L389, A399, D400 and F401 residues (Figure 6(e,f)). However, in contrast to other ABL states, we observed the emergence of additional mediating centers in residues L406, Y408, Y412, L448, I451, T453, Y454 as well as residues from the allosteric binding pocket (Figure 6f)). Notably, the mutation-sensitive allosteric centers also include C-spine residues C388, L389, and I451. The NMR analysis showed a significant role of Y408 in increasing population of the I2 state, and the interactions of the displaced Y412 A-loop residue with the L403/L406/M407 positions are important in allosteric changes of the inactive I2 form.^23^ The structural map of top 20 allosteric centers revealed a narrower and more localized pattern as compared to the active state and I1 (Figure 6(f)). Strikingly, the predicted mediating centers in the inactive I2 state form an allosteric route between the ATP binding site and the allosteric binding pocket, while the long-range communications between the ATP and substrate binding sites become partially depleted.

Our results revealed that the allosteric communications of the ATP site with the substrate binding region and allosteric pockets may coexist and are not mutually exclusive. However, the presented findings indicated a potential shift in the allosteric communication ensemble of the I2 state that favors the long-range couplings between the ATP site and allosteric binding pocket (Figure 6(e,f)). These findings are consistent with the recent NMR studies showing lack of synergy and “weak antagonism” between allosteric inhibitor Asciminib bound in the allosteric site and ATP-competitive inhibitors that stabilize the open αC-helix ABL conformation, whereas in contrast, an ATP-competitive inhibitor of ABL that stabilizes the αC-helix-out, closed conformation showed the increased synergy with Asciminib binding.^97^ According to these experiments, the mechanism by which Asciminib binding inhibits the ABL catalytic function may be related to the synergy between the allosteric site and the ATP site in the inactive ABL conformations.

To summarize, the central finding of the network-based mutational scanning analysis is conformation-specific modulation of the allosteric interactions that may control shifts in the preferential communication routes connecting the ATP site, the substrate binding site and the allosteric binding pocket. Notably, our results and analysis are performed using the experimentally determined conformational states of the unbound ABL catalytic domain and did not consider the complete regulatory ABL-SH2-SH3 construct where the presence of the interacting SH2 and SH3 domains in the assembled state can influence the dynamic properties of the allosteric pocket. Nonetheless, the presented data provided evidence that the allosteric cross-talk between the binding site and regulatory site can be differentially modulated in the active and inactive states through “activation” and “suppression” of the regulatory hotspots and preferential stabilization of conformation-specific communication routes.

### Dimensionality Reduction Analysis and Markov State Model of Allosteric Transitions

While the analysis of MD simulations and functional movements provided important insights into the underlying conformational landscape of ABL, the high dimensionality of the data sets produced by simulations often hinders salient dynamic signatures associated with the mechanisms of allosteric transitions. Here, to facilitate the conformational landscape analysis we employed a deep learning-based dimensionality reduction method to project the results of MD simulations into low dimensional space.^65–67^ The loss function used for the training process in ivis is a triplet loss function that calculates the Euclidean distance among data points and simultaneously minimizes the distances between data of the same labels while maximizing the distances between data of different labels, thus allowing to preserve both local and global structures in a low-dimensional space. The pair-wised Cα distances are often used to characterize protein conformations and movements. In ABL kinase there are 287 residues, leading to 41041 pairs of Cα distances. To obtain a low dimensional space projection of the MD ensembles, ivis dimensionality reduction method was applied to the produced distance dataset. We observed the active state covers a wider region in comparison with the other two states (Figure 7(a)).

**Figure 7.**
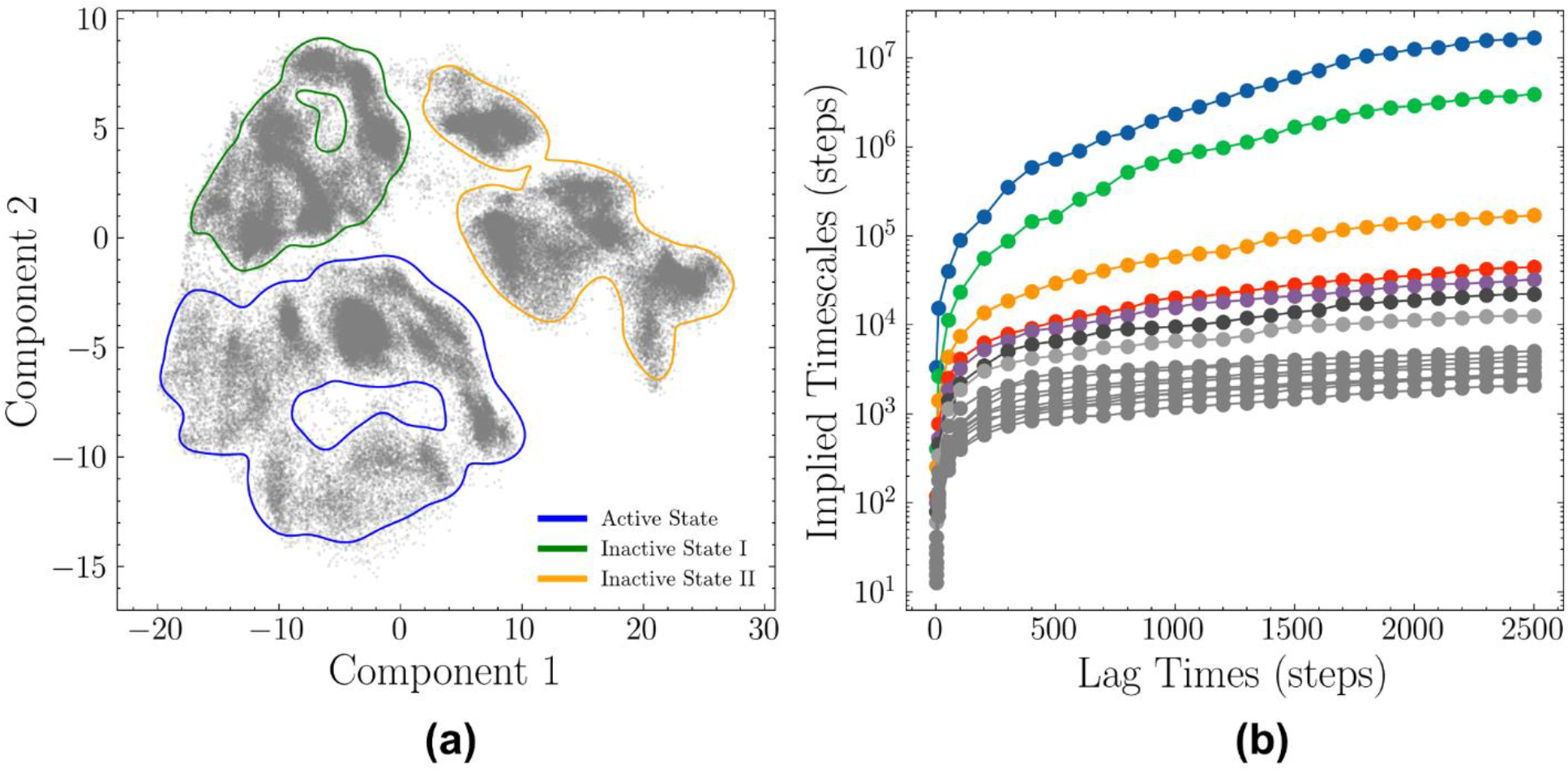
2D ivis dimensionality reduction map and the corresponding implied time scales under different lag times. (A) The distribution of three protein conformations on the reduced 2D space. (B) The estimated relaxation time scale of MSMs under different lag times. The number of steps in MD simulations were used as lag times, ranging from 1 to 2500. In each lag time, an MSM was constructed, and the estimated relaxation time scale was calculated from the transition matrix. The implied time scale converged with a lag time of 2000 steps, which was chosen for MSM construction in further analysis.

Moreover, there is no overlap between the active state and the inactive state I2 but instead a big gap as shown in the ivis 2D surface (Figure 7(a)). In addition, we noticed that the low-dimensionality projection of the MD ensemble highlighted a topological similarity between the active state and the inactive I1 state, while also demonstrating a larger conformational space available for the open active ABL conformation (Figure 7(a)). A different low-dimensional signature was obtained for the inactive I2 conformation, reflecting considerable structural and dynamic rearrangements in this state which are associated with the disassembly and partitioning of the intramolecular interaction networks (Figure 7(a)).

Importantly, compared with dimensionality reduction methods, ivis method was shown to be more robust for constructing MSMs and maintaining high similarity between high and low dimensions with the least information loss. Using the conformational ensembles of the ABL states and ivis dimensionality reduction, we constructed MSM and performed analysis of the macrostates to obtain insights into kinetic aspects of allosteric transformations in ABL. The projected low dimensional space produced by the ivis method (Figure 7(a)) was applied to the MSM construction. In the ivis 2D space, a MiniBatch *k*-means clustering method was applied to partition the 2D data into 500 microstates. The implied timescales were calculated with lag times ranging from 1 to 2500. The top 15 timescales are shown in Figure 7(b). The trend of implied relaxation timescale revealed that the estimated timescale converged after ∼ 2500 steps, which was chosen as the lag time in the construction of MSM. Based on the gap of time scales, the number of macrostates was set to eight and the distribution of macrostates with their transition probabilities enabled analysis of allosteric changes between the active and inactive ABL states (Figure 8). After the partition of the MSM microstates to macrostates was obtained, the stationary distribution and transition probabilities were calculated based on the constructed MSM. According to the MSM partition of states, macrostates 1, 2 and 3 belong to the active kinase conformation, macrostates 4 and 6 are associated with the inactive structure I1 and macrostates 5, 7, and 8 belong to the inactive structure I2 (Figure 8). For each macrostate, the RMSD between its representative structure and the corresponding ABL conformation was calculated. Based on the RMSD values, macrostates 1, 6, and 7 were assigned as the active state, inactive state I, and inactive state I2 respectively, while other five macrostates were designated as intermediate states (Figure 8).

**Figure 8.**
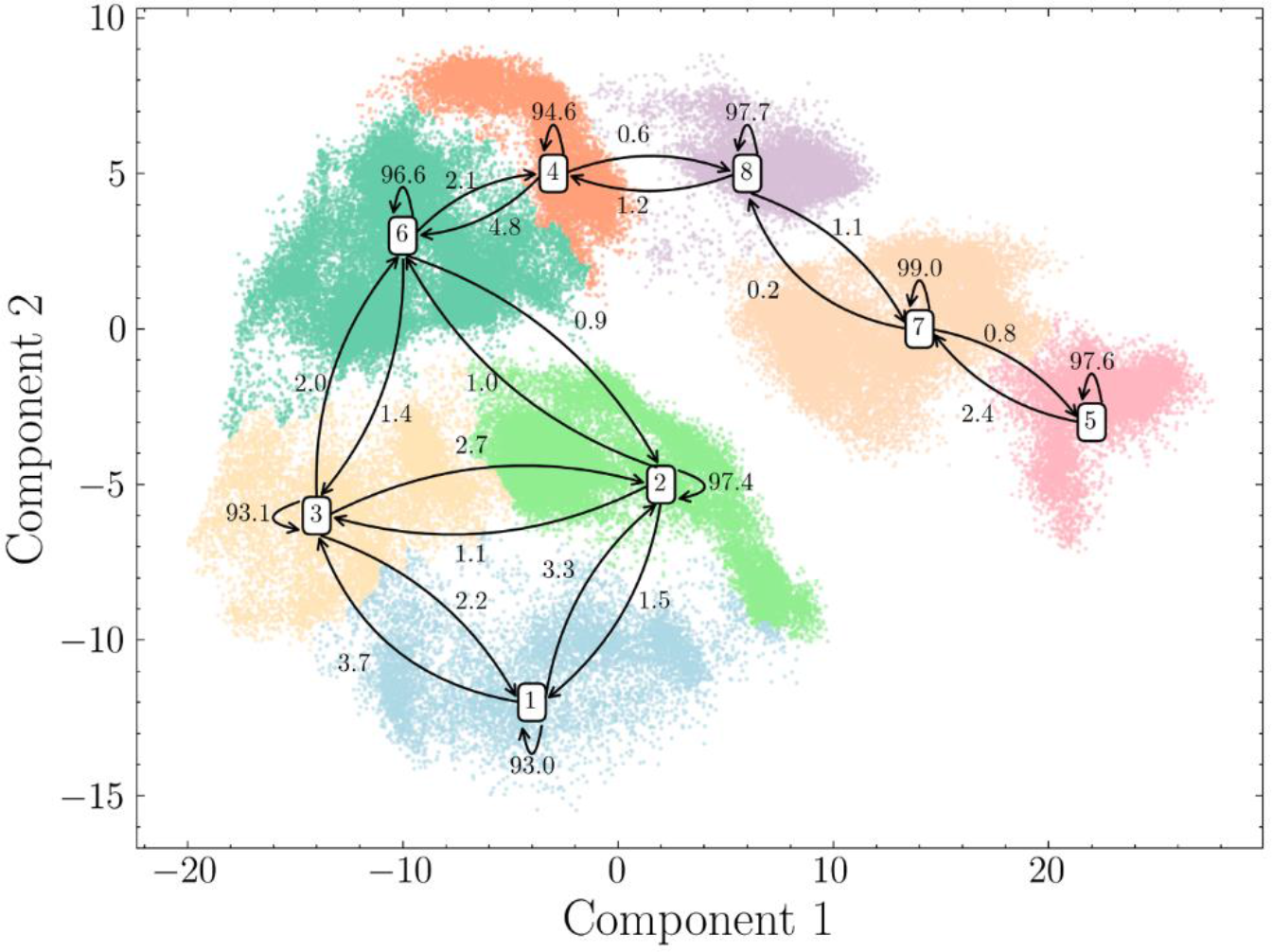
MSM analysis of the ABL conformational landscape. Macrostates 1, 2 and 3 belong to the active kinase conformation, macrostates 4 and 6 are associated with the inactive structure I, while macrostates 5, 7, and 8 belong to the inactive structure I2 . Based on the RMSDs between representative structures in macrostates and the corresponding native structures, macrostate 1, 6, and 7 were considered as the active state, inactive state I1, and inactive state I2, respectively. Other macrostates were treated as intermediate states.

The structural signatures of the obtained ABL macrostates are consistent with the MSM study of the ABL kinase that revealed a total of 16 important macrostates that differed in the functional conformations adopted by the αC-helix (in, out), the DFG-motif (in, out), the P-loop (kinked, extended), and the A-loop (open closed)^31^. We observed that each macrostate is metastable with high probabilities to remain in the same macrostate.

Moreover, the macrostates can more easily interconvert within each of the allosteric ABL states but have higher kinetic barriers for transition to the other states (Figure 8). For example, for the inactive state I1 (macrostates 4 and 6), there is an overall probability of 6.9% to shift between two macrostates, but only 2.3% to transition to the active state and 0.6% to the inactive state I2 (Figure 8). The transition probabilities revealed that ABL kinase can transition following the path from the active state to the inactive state I1 to the inactive state I2 while direct transitions from the active state to the inactive state I2 are kinetically unfavorable. Furthermore, there is an overall 3% probability to transition from the active state to inactive state I1 (2% from macrostate 3 to 6 and 1% from macrostate 2 to 6), but only 0.6% from the inactive state I1 to inactive state I2 (from macrostate 4 to 8). These results are consistent with the experimental findings of a linear equilibrium between the ground active state G and inactive states G↔I1↔I2 where the kinetic exchange between the active and inactive state I2 is slow due to the DFG flip and dramatic rearrangements in the A-loop.^23^

Structural superposition of the microstates 1, 2, and 3 with the structure of the ABL active conformation highlighted the variability of the P-loop and open A-loop (Figure 9A). The P-loop can directly affect ligand binding and experiences transitions from the “kinked” conformation in microstate 1 to an “extended” conformation seen in macrostates 2 and 3 (Figure 9A). Interestingly, the extended P-loop conformation may be a preferable form as it was observed in the large number of protein kinase crystal structures^31^, while a less common partially folded “kinked” conformation is seen in the active ABL form^23^ and is important for selective binding of Imatinib for ABL kinase over c-SRC kinase.^101, 102^ A substantial conformational variability of the open A-loop in the macrostates associated with the active ABL form was also noticeable (Figure 9A). Importantly, in these macrostates, the αC-helix maintains active “in” conformation which is the key signature of the active kinase state (Figure 9A). By aligning the microstates 4 and 6 with the structure of the inactive I1 state, we observed appreciable variations in the main functional regions αC-helix, the P-loop and the A-loop (Figure 9B). While the A-loop in these macrostates undergoes moderate conformational changes, more significant displacements were observed for the αC-helix that experienced movements between the αC helix–in and the inactive αC helix– out positions (Figure 9B).

**Figure 9.**
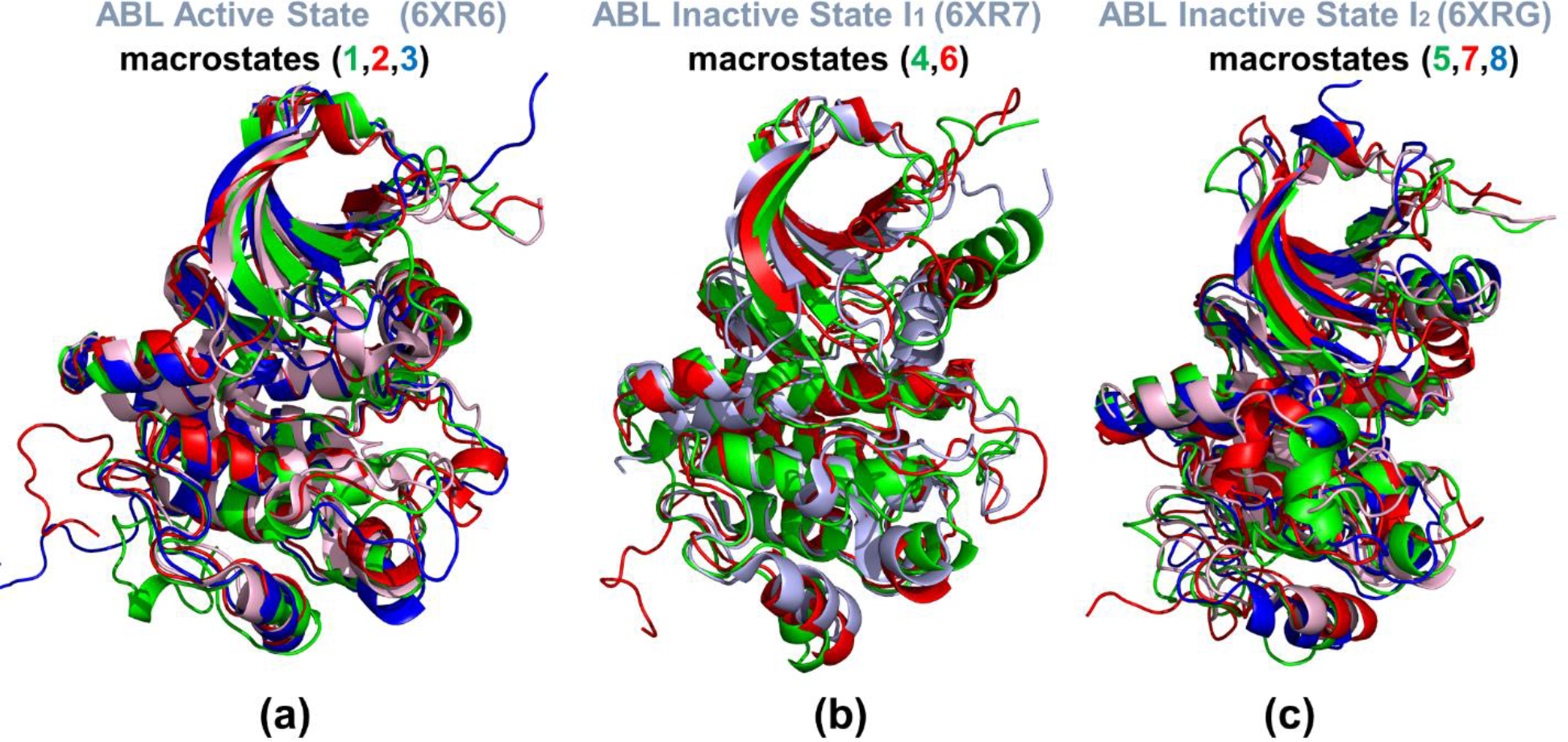
Structural analysis of the ABL macrostates. (a) Structural superposition of the conformations representing macrostates 1(in green ribbons), 2 (in red ribbons) and 3 (in blue ribbons) that belong to the active kinase conformation. The experimental active state conformation is shown in light-pink ribbons. (b) Structural superposition of the conformations representing macrostates 4(in green ribbons) and 6 (in red ribbons) that belong to the inactive I1 state. The experimental I1 conformation is shown in light-pink ribbons. (c) Structural superposition of the conformations representing macrostates 5 (in green ribbons), 7 (in red ribbons) and 8 (in blue ribbons) that belong to the inactive I2 state (shown in light-pink ribbons).

In the I1 structure, the kinase adopts the DFG-out/αC–in conformation and becomes catalytically inactive. The analysis of the I1 macrostates showed that dynamically this inactive state may be susceptible to large movements of the αC-helix that would facilitate conformational transformation to the spectrum of inactive ABL conformations. We also observed large displacements of the “extended” P-loop conformations that are coupled with the reciprocal movements of the αC-helix (Figure 9B). Overall, these findings pointed to the globally increased functional movements of the key functional regions in the I1 macrostates .

Structural variations among microstates 5, 7, and 8 that belong to the inactive I2 state are even more significant particularly in the N-lobe and A-loop (Figure 9C). We noticed appreciable displacements of the P-loop that can adopt various “extended” stretched conformations that are necessary to accommodate diverse closed conformations of the A-loop. At the same time, the αC- helix movements are moderate and similar to the αC-helix-out position in the inactive I2 structure (Figure 9C). The macrostates revealed a remarkable diversity of the closed A-loop conformations that may be available in the I2 state. It was proposed in the original structural study that Imatinib could bind preferentially to the ABL I2 state but induces additional changes to optimize the binding energy.^23^ MSM analysis indicated that Imatinib binding can induce a less intrinsically favorable “kinked” conformation of the P-loop and significantly reduce mobility of the closed A-loop. As a result, Imatinib binding may significantly reduce conformational mobility and impair favorable entropy intrinsically present in the I2 state that could only be partly compensated by the ligand-protein interactions and result in the observed moderate binding affinity.^23^ The structural analysis of the macrostates associated with the low-populated forms of the ABL kinase can expand our understanding of “hidden” allosteric states and invisible aspects of protein kinase functionalities. While computationally predicted inactive macrostates represent short-lived intermediates, their dynamic proximity to the experimentally determined inactive conformations may be leveraged for the design of selective inhibitors with reduced sensitivity to drug-resistant mutations.

In addition, we also applied a supervised machine learning model, XGBoost^88^ to extract key functional residues that are important in the ABL conformational transitions. The pair-wised Cα distances were used as features and the macrostate numbers were used as labels. The XGBoost model was able to differentiate Abl macrostates with a test accuracy of 99.4% with 300 base estimators and maximum tree depth of 4. The feature importance of Cα distances were calculated. For each distance, the corresponding importance was further accumulated to the related two residues, in which the importance at residue level can be quantified. The top 10 residues are highlighted in Supporting Information Figure S1A as M407, D440, T408, D400, Y412, D523, V299, A418, I379, and E481 in descending order. According to the NMR analysis^23^ half of the identified 10 residues lie in the A-loop, which is vital between the conformation transitions from the active state to the inactive states. The active state structure was further analyzed through Protein Allosteric Sites Server (PASSer).^103^ PASSer is a machine learning based web server for accurate protein allosteric sites prediction. Through the default ensemble model, two promising allosteric sites can be identified (Supporting Information Figure S1B). These two pockets are predicted as the top 2 pockets with probabilities as 36.4% and 34.7%, respectively. One pocket is located in the P-loop and the other one in the A-loop. Both loops undergo significant conformational changes in the ABL protein. Together with the currently validated binding sites, the identified pockets may play a relevant role in mediating allosteric communications and could become “activated” in certain hidden conformational states of ABL. Functional probing the intramolecular communication networks in the ABL conformations and potential allosteric pockets through mutations of the predicted functional positions and vulnerable network links may be useful for engineering and modulating kinase activities.

## Conclusions

In this work, we introduced and synergistically applied several computational approaches to examine at atomistic details conformational landscapes and mechanisms of allosteric regulation in the distinct functional states of the ABL kinase domain. By using MD simulations and ensemble-based distance fluctuations analysis, we characterized a critical role of dynamic coupling between N387 residue, the regulatory 400-DFG-402 motif and the R-spine residues in mediating structural stability and allosteric communications in the ABL states. Our results revealed a previously unappreciated role of N387 as a regulatory anchor that couples functional kinase regions to modulate allostery in the kinase domain. These results are consistent with the recent biophysical experiments and explain the importance of N387 in modulating long-range functional effects in the ABL kinase.

We proposed a network-based mutational profiling approach for probing of allosteric residue propensities and communications that revealed a group of conserved allosteric regulatory hotspots in the ABL states. The results of mutational profiling modulation of the allosteric interaction networks between the active and inactive states where the long-range communication pathways between the ATP binding site and substrate regions are dominant in the active state but are partially suppressed in the closed inactive state I2 and replaced with the stronger allosteric couplings are between the ATP site and the allosteric binding pocket. The important finding of this study is evidence of conformation-specific modulation of the allosteric interaction networks and communications that can alter preferential communication routes and cross-talk between the ATP site, the substrate binding site and allosteric site. These results agree with the latest NMR experiments suggesting a stronger cooperativity between the ATP-binding site and allosteric pocket in the inactive closed conformation.^97^

By integrating the results of MD simulations with MSM-based identification of macrostates we analyzed the kinetic transitions between the ABL functional conformations that follow the path from the active state to the inactive state I and then to the inactive state I2, while direct transitions from active state to inactive state I2 are not favorable. Through structural analysis of the ABL macrostates, we showed that conformational variability and long-range cooperativity between the functional regions of αC-helix, the P-loop and the A-loop can provide a concerted mechanism for allosteric transitions underlying regulation of the kinase activity. We argue that probing and exploiting diverse ensemble of the intramolecular communication networks in the ABL conformations through targeted modifications of vulnerable network links and inter-community bridges may be useful for engineering and modulating kinase activities. Together with NMR studies of kinase dynamics and advances in deep mutational scanning of allostery^104–106^ the proposed computational approach may open up new venues for probing kinase-centric signaling processes and engineering of allosteric functions.

## Acknowledgment

GV acknowledges support by the Kay Family Foundation Grant A20-0032.

## CRediT author statement

**Keerthi Krishnan:** Data curation, Methodology, Software, Validation, Visualization, Writing- Original draft preparation **Hao Tian.**: Data curation, Methodology, Validation, Visualization, Software Writing- Original draft preparation. **Peng Tao**: Data curation, Methodology, Validation, Visualization, Software Writing- Original draft preparation. Investigation. **Gennady Verkhivker**: Conceptualization, Supervision, Data curation, Methodology, Software, Validation, Visualization, Validation, Writing- Original draft preparation, Writing- Reviewing and Editing

## Data Availability Statement

The data that support the findings of this study are available from the corresponding author upon reasonable request.

## No Conflicts of Interest

The authors have no conflicts to disclose.

The authors declare that the research was conducted in the absence of any commercial or financial relationship that could be construed as a potential conflict of interest. The funders had no role in the design of the study; in the collection, analyses, or interpretation of data; in the writing of the manuscript, or in the decision to publish the results.

