## Supporting Figure S1 for "Probing Conformational Landscapes and Mechanisms of Allosteric Communication in the Functional States of the ABL Kinase Domain Using Multiscale Simulations and Network-Based Mutational Profiling of Allosteric Residue Potentials"

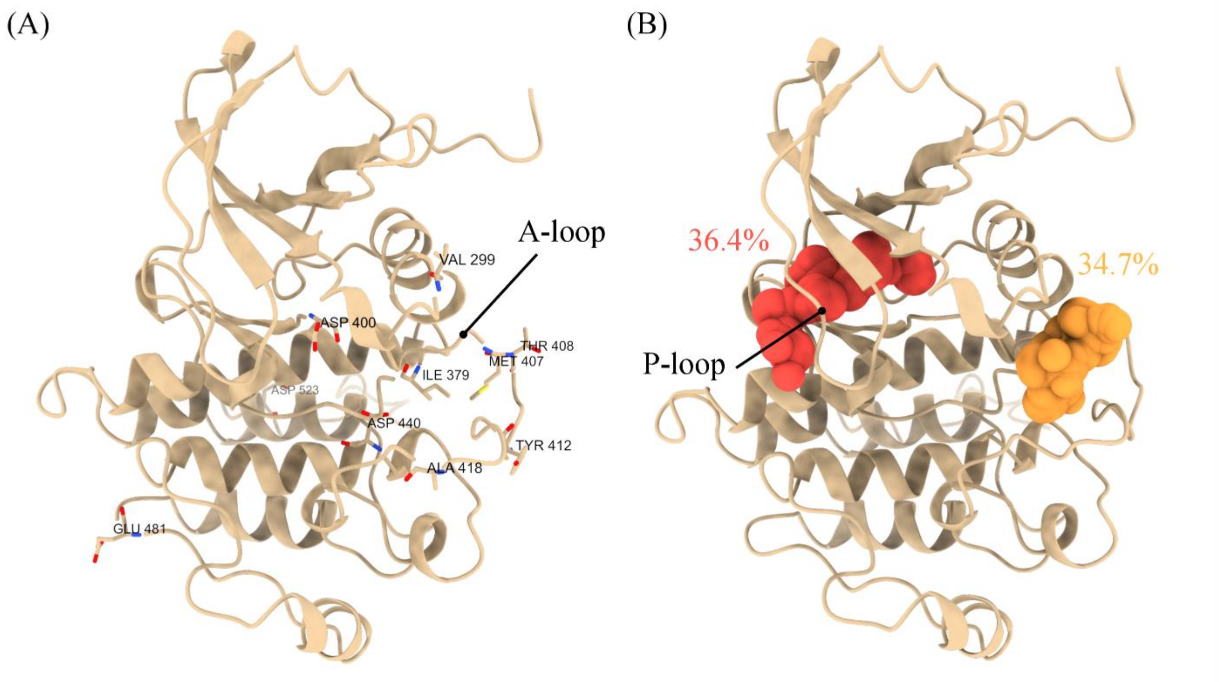


**Figure S1.** Machine learning identified important residues and pockets. (A) The most important 10 residues are highlighted as M407, D440, T408, D400, Y412, D523, V299, A418, I379, and E481 in descending order. Half of them lie in the A-loop, which is important in initiating the transition from active to inactive states. (B) The top 2 protein pockets identified by PASSer. One is located in the P-loop region, and another is in the A-loop region, and both with high probabilities as the potential allosteric sites.
